# Chrom-Sig: de-noising 1-dimensional genomic profiles by signal processing methods

**DOI:** 10.1101/2025.08.12.670000

**Authors:** Nandita J. Gupta, Zachary Apell, Minji Kim

**Affiliations:** Department of Electrical and Computer Engineering, University of Michigan, Ann Arbor, MI, USA; Gilbert S. Omenn Department of Computational Medicine and Bioinformatics, University of Michigan, Ann Arbor, MI, USA; Department of Biostatistics, University of Michigan, Ann Arbor, MI, USA

## Abstract

**Motivation:** Modern genomic research is driven by next-generation sequencing experiments such as ChIP-seq, CUT&Tag, and CUT&RUN that generate coverage files for transcription factor binding, as well as ATAC-seq that yield coverage files for chromatin accessibility. Due to the inherent technical noise present in the experimental protocols, researchers need statistically rigorous and computational efficient methods to extract true biological signal from a mixture of signal and noise. However, existing approaches are often computationally demanding or require input or spike-in controls.

**Results:** We developed Chrom-Sig, a Python package to quickly de-noise 1-dimensional genomic coverage tracks by computing the empirical null distribution without prior assumptions or experimental controls. When tested on 19 ChIP-seq, CUT&RUN, ATAC-seq, snATAC-seq datasets, Chrom-Sig can effectively decompose the data into signal and noise. Notably, Chrom-Sig performs de-noising and peak calling in 1-2 hours using around 20 GB of memory. The de-noised signal corroborates with biologically meaningful results: CTCF CUT&RUN data retained a higher percentage of peaks overlapping CTCF binding motifs and ATAC-seq and RNA Polymerase II data resulted in enhancers and promoters. We envision Chrom-Sig to be a highly versatile and general tool for current and future genomic technologies.

**Availability:** Chrom-Sig is publicly available at https://github.com/minjikimlab/chromsig under the MIT license.

## 1 Introduction

The advances in high-throughput sequencing technologies have enabled researchers to probe various activities and structures in the genome. For example, one can probe gene expression (RNA-seq) (Mortazavi *et al*., 2008), protein binding intensity or histone modification (ChIP-seq, CUT&RUN, CUT&Tag) (Robertson *et al*., 2007; Skene and Henikoff, 2017; Kaya-Okur *et al*., 2019), chromatin accessibility (ATAC-seq) (Buenrostro *et al*., 2015), and chromosome conformation (Hi-C, ChIA-PET) (Lieberman-Aiden *et al*., 2009; Fullwood *et al*., 2009). More recently, single-cell and multi-ome methods have provided the unprecedented resolution and information to simultaneously probe multiple modalities in a single-cell (snATAC-seq, HiRES) (Preissl *et al*., 2018; Liu *et al*., 2023). Due to the significance of genome function and structures on human health, the National Institutes of Health ENCODE (ENCOD Project Consortium, 2012) and 4D Nucleome (Dekker *et al*., 2017; Reiff *et al*., 2022) consortia have led concerted efforts to generate the genomic data in diverse mammalian tissues and cell types.

While these methods have the potential to uncover meaningful biological phenomena, the progress is often impeded by the inherent technical bias and noise present in the experimental protocols. To avoid misinterpreting the data, one needs to apply a rigorous statistical method to assess the significance of observed data compared to the background null model. However, new assays often lack a known theoretical background distribution. One approach to mitigate the problem is to perform input control or spike-in experiments to obtain the experimental background, but doing so can be laborious and costly or infeasible for rare samples.

As an alternative solution, researchers have developed computational algorithms Coda (Koh *et al*., 2017) and AtacWorks (Lal *et al*., 2021) to de-noise 1-dimensional ChIP-seq or ATAC-seq data, respectively. However, these methods are based on deep-learning and can be computationally intensive, require a large training set that may be lacking for new assays, or overfit the data. An ideal method should be computationally efficient, statistically rigorous, and biologically interpretable—all without requiring additional wet-lab experiments.

Towards this goal, we developed a Python package Chrom-Sig, which quickly obtains empirical background distribution to assign a statistical significance to each observed read, thereby retaining reads with high significance as means to extract true biological signal in any 1-dimensional genomic assays.

## 2 Methods

A simple, yet powerful way to probe the empirical null distribution is to place each observed read in a random location within the same chromosome many times and recording the signal intensity therein. However, the key bottleneck of this approach is the efficiency: searching for a given interval in a large bedGraph file is a computationally expensive job. This problem is exacerbated by the fact that robust statistics requires a large number of samples. To overcome this problem, we have previously developed pyBedGraph (Zhang *et al*., 2020), which can quickly obtain summary statistic from 1-dimensional genomic signal. Specifically, obtaining the exact mean for 10 billion intervals is estimated to take 43 minutes with pyBedGraph and 7.4 days with a similar software pyBigWig (Ramírez *et al*., 2016).

By leveraging pyBedGraph, we implemented the enrichment test—assessing the statistical significance of each observed read by comparing the observed signal to the empirical null distribution in a random location—in a Python package Chrom-Sig (**Figure S1**). As an input, Chrom-Sig accepts BAM or bed files of aligned paired-end or single-end reads generated by any 1-dimensional genomic assays including ChIP-seq, CUT&RUN, ATAC-seq, and snATAC-seq. The enrichment test is performed on the coverage bedGraph file generated by piling up reads, resulting in the ‘pass’ or ‘fail’ reads based on the false discovery rate (FDR) given the number of pseudo-reads for the empirical null. The pile-up of ‘pass’ reads further undergo a peak calling algorithm SICER (Xu *et al*., 2014), providing an accurate set of peaks for users. Chrom-Sig is implemented in Python3 using a few strategies to further optimize speed and memory usage. Detailed methods are provided in **Methods S1, S2**.

## 3 Results

We applied Chrom-Sig on 11 paired-end and 8 single-end datasets (**Table S1**) from ChIP-seq, CUT&RUN, ATAC-seq, and snATAC-seq experiments in GM12878 and K562 cell line downloaded from the ENCODE portal (Sloan *et al*., 2016) (https://www.encodeproject.org) and 4DN data portal (Dekker et al., 2017; Reiff et al., 2022). All runs were on an Intel® Xeon Gold 6154 CPU @ 3.0 GHz.

We first ran Chrom-Sig with 5000 pseudo-reads (**Table S2**). The paired-end data consisted of 7-174 million uniquely mapped reads and FDR of 0.2 retained 11-76% of them, while FDR of 0.1 kept 5-67% (**Figure S2a**). For single-end data, there were between 4-15 million uniquely mapped reads, of which 1.5-35% and 0.5-30% were kept for FDR of 0.2 and 0.1, respectively (**Figure S2b**). Doing so took between 0.06-2.9 hours and 2-85 GB of memory (**Figure S3**). To evaluate the effect of the number of pseudo-reads on the runtime and % of pass, we varied the pseudo-reads from 50 to 5000 on GM12878 CTCF ChIP-seq and RAD51 ChIP-seq data. As expected, the runtime had a linear trend with ∼100 seconds on 1000 pseudo-reads and ∼400 seconds on 500 pseudo-reads, and the % of pass reads converged to 6.8% for paired-end and 14 % for single-end after 2500 pseudo-reads (**Figure S4**).

For example, the GM12878 CTCF ChIP-seq data were originally noisy with a large portion of reads in non-binding sites, but Chrom-Sig with FDR of 0.1 retained only the reads with strong binding (**Figure 1a**). Similarly, CTCF CUT&RUN data originally had a high number of false positive peaks due to noise. After running Chrom-Sig with FDR of 0.2 or a more stringent 0.1, the peaks identified by SICER on Chrom-Sig ‘pass’ (signal) were highly specific; the ‘fail’ (noise) pile-up is non-specific and resembles the data from input control experiments (**Figure 1a**). This general trend is observed for all 19 datasets, where the de-noised data are visually cleaner and the peaks contain less false positives (**Figures S5, S6, S7**). In particular, the two YY1 ChIP-seq data were highly noisy and of low-depth (∼5 million reads), yet Chrom-Sig was able to recover a small number of peaks that coincide with those of two other replicates with high-depth (∼15 million reads) (**Figure S7**).

**Fig. 1.**
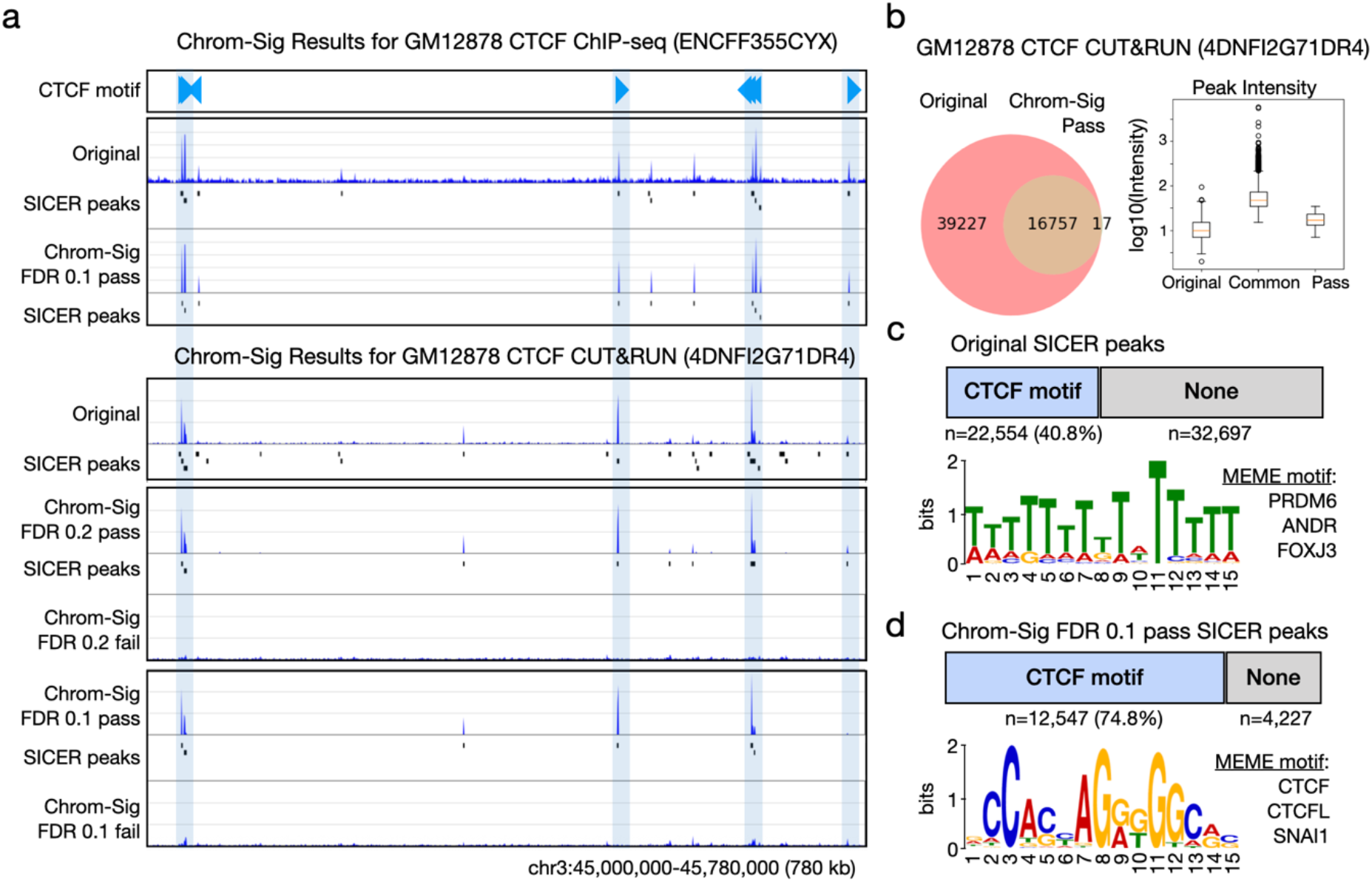
Chrom-Sig results on GM12878 CTCF ChIP-seq and CUT&RUN datasets. a) Browser views of CTCF binding motifs with orientation (blue triangles) and coverage tracks generated by piling up the original (before Chrom-Sig) and Chrom-Sig ‘pass’ or ‘fail’ reads, accompanied by the peaks called by SICER. b) Venn Diagram of the peaks called on the original and Chrom-Sig FDR 0.1, 5000 pseudo-reads ‘pass’ reads pile-up, with boxplots of maximum peak intensity for each peak. c) Number of peaks overlapping CTCF motifs, and top MEME result on the original data. d) Similar to panel c for Chrom-Sig ‘pass’ results.

To compare the peaks before and after Chrom-Sig, we computed the proportion of overlap. As visually speculated for GM12878 CTCF CUT&RUN data (4DNFI2G71DR4), there were originally 55,251 peaks, of which 16,757 (42.7%) were recovered by Chrom-Sig; it also identified a small number (17) of new peaks (**Figure 1b**). The binding intensity for each group of peaks verified that the common peaks (‘Common’) were the strongest—likely representing true CTCF binding sites—followed by Chrom-Sig-specific (‘pass’) and Original-specific (‘Original’) groups (**Figure 1b**). Similar results are shown for all 19 datasets (**Figure S8, S9**).

Finally, we sought to verify that Chrom-Sig retains biologically meaningful signal while removing false positive peaks. Using the known CTCF motif binding sites as a feature of true signal, we verified that the original SICER peaks had only a small proportion (40.8%) with the top MEME (Machanick and Bailey, 2011) motif result showing the enrichment of PRDM6, ANDR, FOXJ3 instead of CTCF (ranked 18) (**Figure 1c**). By contrast, Chrom-Sig peaks mainly overlapped CTCF motifs (74.8%) and were strongly enriched in CTCF motifs via MEME (**Figure 1d**). Top 3 MEME motifs are also presented (**Figure S10a**). We verified with other GM12878 CTCF ChIP-seq and CUT&RUN data that Chrom-Sig was more specific in identifying peaks overlapping known CTCF motifs (**Figure S10b**). Similarly, Chrom-Sig specifically enriched for active promoters, enhancers, and transcription in ATAC-seq and RNA Polymerase II ChIP-seq data—as expected and desired (**Figure S11**).

## 4 Discussion

We developed Chrom-Sig and demonstrated its ability to de-compose 1-dimensional genomic data into signal (‘pass’) and noise (‘fail’). Specifically, de-noising the data with Chrom-Sig resulted in highly specific peaks that correlated with biological evidence such as CTCF binding motifs and active chromatin states. However, one drawback of the current version is the lack of guidance on FDR threshold and the number of pseudo-reads. We generally recommend FDR of 0.2 and 5000 pseudo-reads, as a stringent FDR may unintentionally remove signal and low pseudo-reads may not provide enough statistical power. In addition, we intend to compare Chrom-Sig to deep-learning-based methods such as Coda and AtacWorks in the future. As the field of genomics continue to develop novel experimental technologies to measure multiple types of biological signal at single-cell level, it will be imperative to distinguish true signal from technical noise. We envision Chrom-Sig to be instrumental in filtering out inherent technical noise without requiring input or spike-in control experiments, thereby providing a versatile and widely applicable tool for researchers.

## Public data

Authors thank members of the ENCODE and 4DN consortia for generating and depositing the data, including the GM12878 CUT&RUN data from the Henikoff lab.

## Supplementary Method S1. Chrom-Sig algorithm

For each dataset to be processed, Chrom-Sig takes either a bam file or bed file as input. Bam or bed file inputs often contain non-standard chromosomes, which significantly slow down the process of generating a bedgraph file. Chrom-Sig therefore uses the reference genome sizes file as indication for which chromosomes to keep, and generates a new standard bed file, containing only the standard chromosome data from the bed file. Converting this standard bed into a bedgraph is as much as 30x faster than converting from the original bed file.

Since each chromosome must be processed separately when de-noising, Chrom-Sig generates a separate node for each chromosome in order to run all chromosomes in parallel. Not all datasets contain all chromosomes – Chrom-Sig generates nodes for the maximum number of chromosomes that could be contained in the data. Within each node, the denoising process generates a list for which chromosomes are in the bed file, and only runs the Python program for de-noising on those chromosomes. During this process of checking which chromosomes have data, it also checks for the coverage of each chromosome. If the data in the bed file has less than 20% coverage for a given chromosome, then that specific chromosome is not processed. In de-noising each chromosome separately, only the data for the chromosome in hand needs to be stored. The first step in the de-noising process is to read in the bed file – instead of reading in all the data, Chrom-Sig only reads in the data for the current chromosome, ignoring the remainder of the bed file.

The most computationally expensive part of Chrom-Sig’s denoising algorithm is in generating and processing pseudo-reads. Pseudo-reads are used to determine expected enrichment values with which to compare the observed samples in the bed file. Since paired-end reads datasets have more variation in read lengths and have the added complication of multiple fragments per read, Chrom-Sig analyzes single-end and paired-end reads datasets separately as follows.

One single-end dataset has the same read length for all reads. Thus, as pseudo-reads for a single read are translations of that read to random locations in the chromosome, Chrom-Sig needs to generate only a single set of pseudo samples to compare all the observed samples with. This strategy significantly reduces the runtime, since for an input x number of pseudo-reads, instead of processing x samples per read, Chrom-Sig processes x samples in total.

A paired-end dataset contains reads which vary in distance between fragments, and the lengths of each fragment. Therefore, distinct variations in fragment length and read span need a unique set of pseudo samples to compare to. While there is significant variation in a paired-end dataset, there is also enough repetition in fragment and read lengths such that, by grouping together samples which share fragment lengths and spans and generating a single set of pseudo samples for each grouping, Chrom-Sig is able to significantly reduce the necessary computation as it did for the single-end datasets.

Determining the enrichment value for a paired-end dataset requires finding the maximum along both fragments in a read, and taking the average of the maximums. Determining this maximum requires calling the pyBedGraph module. To reduce the number of times the module is called as means to reduce the runtime, instead of determining the maximums and calculating the enrichment for each pseudo sample separately, Chrom-Sig creates a list of all fragments for all pseudo-reads, without grouping fragments by which read they belong to. Since lists maintain order, every two fragments in this list belong to a single read. pyBedGraph is then applied to the whole list. Therefore, for a single span, instead of calling pyBedGraph the same number of times as the number of pseudo-reads, it only needs to be called once. From there, pyBedGraph produces a list of maximums for all the fragments, and Chrom-Sig averages every two elements of this list to produce a list of expected enrichments.

The de-noising step that takes the longest time is the process of determining the raw p-values of the observed samples. Each observed enrichment is compared to its corresponding set of expected enrichments, and the number of enrichments for which the observed value is less than the expected is totaled and divided by the number of pseudo samples to determine the raw p-value. While forming a Boolean array for which expected enrichment values were greater than the observed and taking the sum of this array may take fewer lines of code, taking the sum of an array is computationally expensive. Instead, Chrom-Sig iterates through each expected enrichment value, comparing it to the observed, and adds the result to a running total. It then divides this total by the number of pseudo samples to optimize the speed.

Chrom-Sig outputs the passing and failing samples into separate beds. In order to visualize the results, Chrom-Sig then converts these bed files into bedgraph files. This process also converts the ‘pass’ and ‘fail’ beds to bedgraphs for all chromosomes in parallel. Finally, when each chromosome-wise processes are completed, Chrom-Sig concatenates all the pass beds, fail beds, pass bedgraphs, and fail bedgraphs into “total” files for the whole-genome results.

Chrom-Sig also produces peak-calling results for both the original input data and the Chrom-Sig results. It runs the SICER2 algorithm, which takes a bed file and the name of the reference genome as inputs, and outputs a scoreisland file with the locations of the peaks it called. Chrom-Sig runs SICER2 on the original bedgraph file, and the total pass-pileup bedgraph file.

## Supplementary Method S2. Chrom-Sig usage and commands

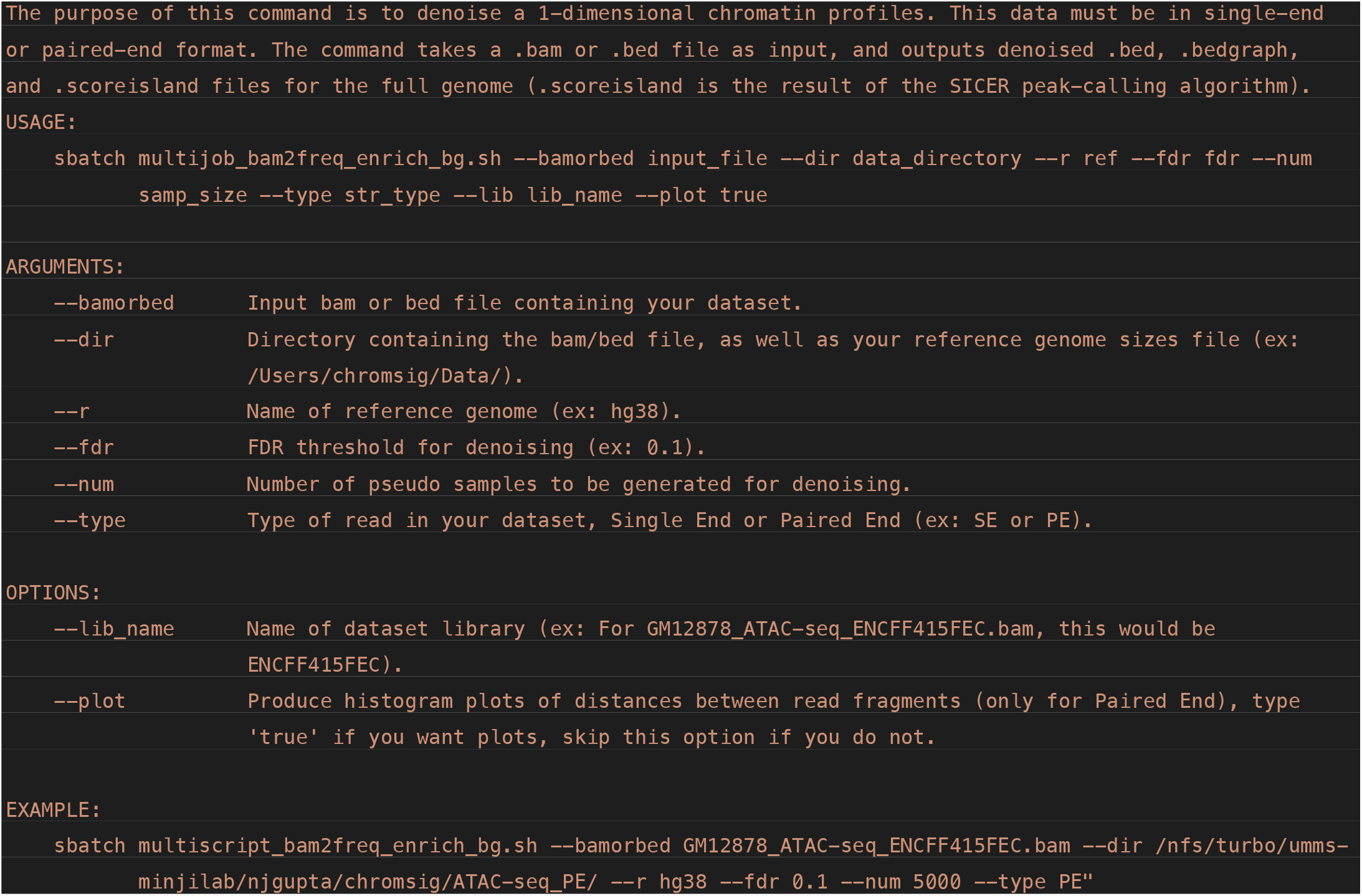

## Supplementary Method S3. Analysis of Chrom-Sig results

### CTCF Motifs

To assess the precision of CTCF binding site identification before and after Chrom-Sig with FDR 0.1 and 5000 pseudo-reads, we implemented a python script using python 3.11.11 with matplotlib version 3.9.2, pybedtools 0.10.0, and numpy 2.2.1. The analysis looks at GM12878 CTCF ChIP-seq ENCFF355CYX (36,269 peaks original, 24,872 peaks after Chrom-Sig) as well as GM12878 CTCF CUT&RUN replicates 4DNFI2G71DR4 (55,251 peaks original, 22,554 peaks after Chrom-Sig) and 4DNFI9U71IB4 (62,176 peaks original, 19,233 peaks after Chrom-Sig).

CTCF motif locations were provided in bed format called by STORM as described in Tang et al., 2015. Peak files were provided in a bed format. Genomic intervals were loaded using pybedtools and the number of regions in each file were counted. We defined precision as the overlap of peaks with CTCF motifs divided by the number of total peaks. Overlap between CTCF motifs and peak regions was calculated using pybedtools intersect with parameter u=True to obtain the number of CTCF motif regions in each peak file. Both the Chrom-Sig pass pileup SICER peak file and the SICER peak calls on the original data were analyzed for each sample.

### MEME Analysis

In continuity with the above analysis, motif analysis was performed on GM12878 CTCF CUT&RUN 4DNFI2G71DR4 for the original data and after Chrom-Sig with FDR 0.1 and 5000 pseudo-reads. The MEME suite (v5.5.5) was used to identify enriched DNA sequence motifs within peak regions using the command line version. The input required for this process is peak region file in a bed format, a reference genome, and motif database. We used bedtools version 2.31.1 to convert to fasta through the getfasta command with hg38 reference genome from iGenomes/Homo_sapiens/UCSC/hg38/Sequence/BWAIndex/genome.fa. The motif database HUMAN/HOCOMOCOv11_core_HUMAN_mono_meme_format.meme was downloaded from MEME. The meme-chip tool was run with default parameters (-time 240 -ccut 100 -dna -order 2 -minw 6 -maxw 15 -db HUMAN/HOCOMOCOv11_core_HUMAN_mono_meme_format.meme -meme-mod zoops -meme-nmotifs 5 -meme-searchsize 100000 -streme-pvt 0.05 -streme-align center -streme-totallength 4000000 -centrimo-score 5.0 -centrimo-ethresh 10.0) on each fasta file. The output was an html meme report with motifs, E-value, discovery program, and known motif matches from the database. We reported at the top 3 motifs for both before (‘original’) and after Chrom-Sig (‘pass’). The script “run_meme.sh” was run on a cluster with 10G of memory and took less than an hour to run for each peak file.

### ChromHMM States

This analysis compares the distribution of ChromHMM states in original and Chrom-Sig processed datasets across GM12878 ENCFF646NWY ATAC-seq data and K562 RNAPII ChIP-seq replicates ENCFF480AJZ and ENCFF785OCU. ATAC-seq peaks and RNAPII ChIP-seq peaks are known to be correlated with promotors and enhancers and our aim was to evaluate whether Chrom-Sig enhances such known correlations.

A script named “chromHMM_annotation.sh” took as input previously annotated region files for GM12878 and K562 cell lines and the aforementioned peak files for both original data and Chrom-Sig. The script used bedtools 2.31.1 intersect with option -wao to get the size of the intersection where -a is the peak file and -b is the annotation file. The output was saved as the peak file name with “_chromhmmanno.bed” extension.

A custom python workflow named “chromsig_chromhmm_annotation.ipynb” takes the annotated files as input and outputs the plots in the manuscript. The method uses Python 3.9.23 and packages numpy and matplotlib. ChromHMM states are grouped to reduce the state space from 15 to 7. The 7 groups are promoter, enhancer, transcription, insulator, repressed polycomb, heterochromatin, or other. The percentage of each state is calculated by summing the number of intersecting base pairs for each state and dividing by the total base pairs. These percentages were calculated for both original and Chrom-Sig processed datasets and are shown as stacked bar charts.

**Supplementary Figure S1.**
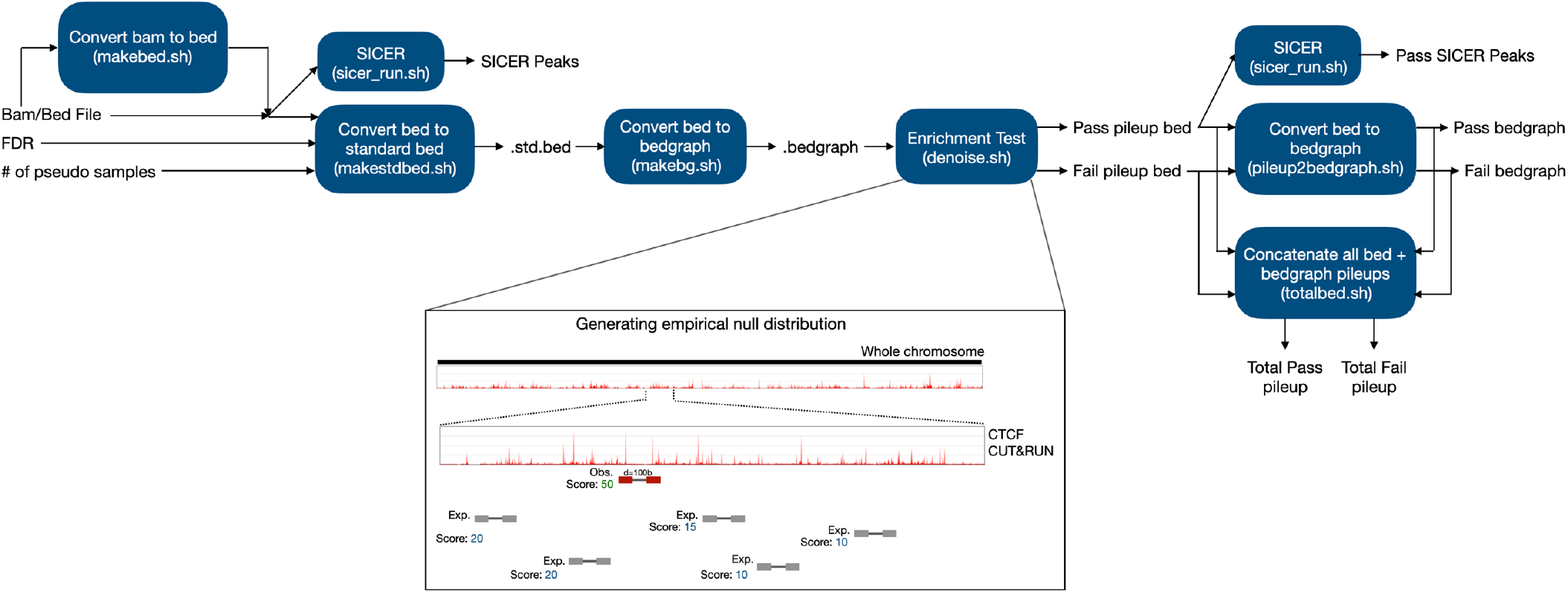
Overview of `Chrom-Sig` Chrom-Sig pipeline. If taking a BAM file as input, converts it to a bed file, otherwise takes a bed file as input. FDR: False Discovery Rate threshold. # of pseudo samples: the number of pseudo-reads for the Enrichment Test (denoise.sh) to generate. The pipeline converts a bed file to a standard bed file by sorting the data and removing non-standard chromosomes. The standard bed file is converted to a bedgraph, which is then input along with the bed file to the Enrichment Test. This test generates a number of pseudo-reads specified by the user, determines the maximum value in the bedgraph along each of those pseudo-reads (for paired end data, taking the average of the maximums along each fragment), and compares the maximum along the observed read (from the bed file) to the pseudo-reads to generate a p-value. If the p-value is below the FDR threshold, the read passes and is copied to a pass-pileup bed file, otherwise it is copied to a fail-pileup bed file. Chrom-Sig then coverts the pass and fail bed files for each chromosome to bedgraph files, and concatenates each set of files into a total file for the whole genome. The SICER peak-calling algorithm is then run on the initial bed file and the total pass-pileup bed file.

**Supplementary Figure S2.**
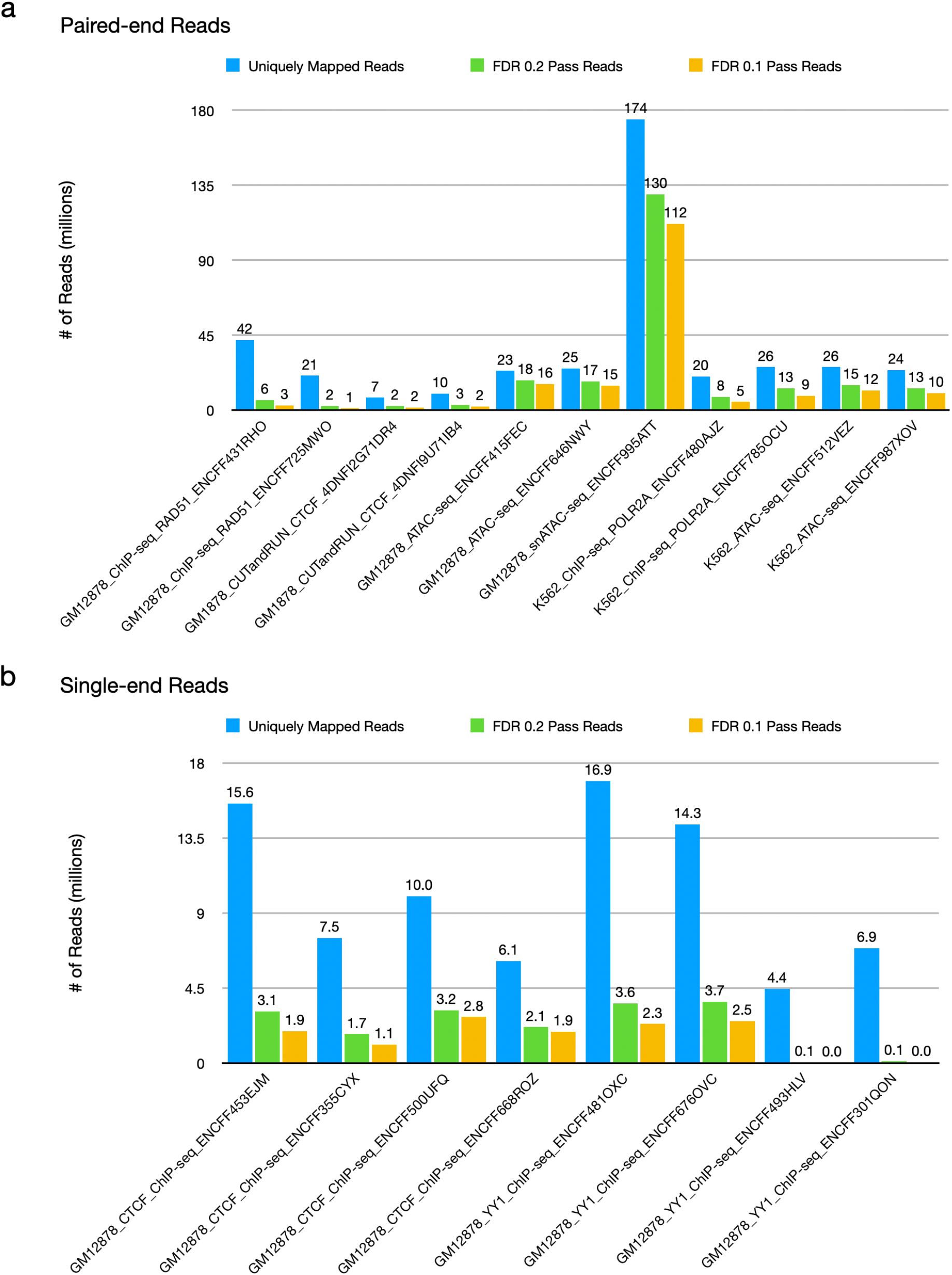
Overview of Datasets Uniquely mapped reads, number of reads retained for FDR threshold of 0.1, and number of reads retained for FDR threshold of 0.2. Uniquely mapped reads were obtained from the original bed file for each dataset. The number of reads is counted in millions. a) 11 paired-end datasets. b) 8 single-end datasets.

**Supplementary Figure S3.**
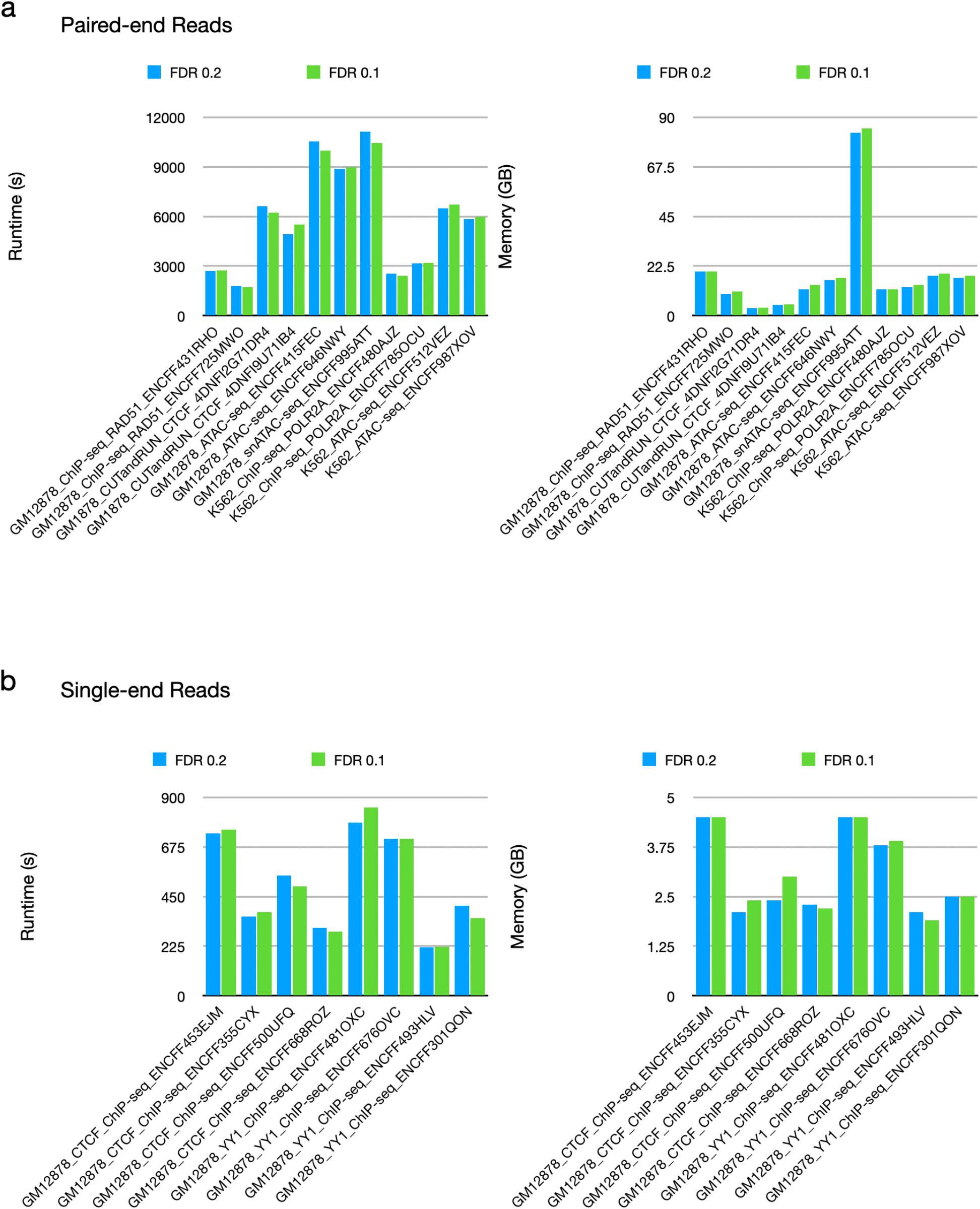
Runtime and Memory Runtime and memory usage for FDR thresholds of 0.2 and 0.1. All runs were done with 5000 samples. Runtime is in seconds, and memory usage is in gigabytes. a) paired-end reads data, b) single-end reads data.

**Supplementary Figure S4.**
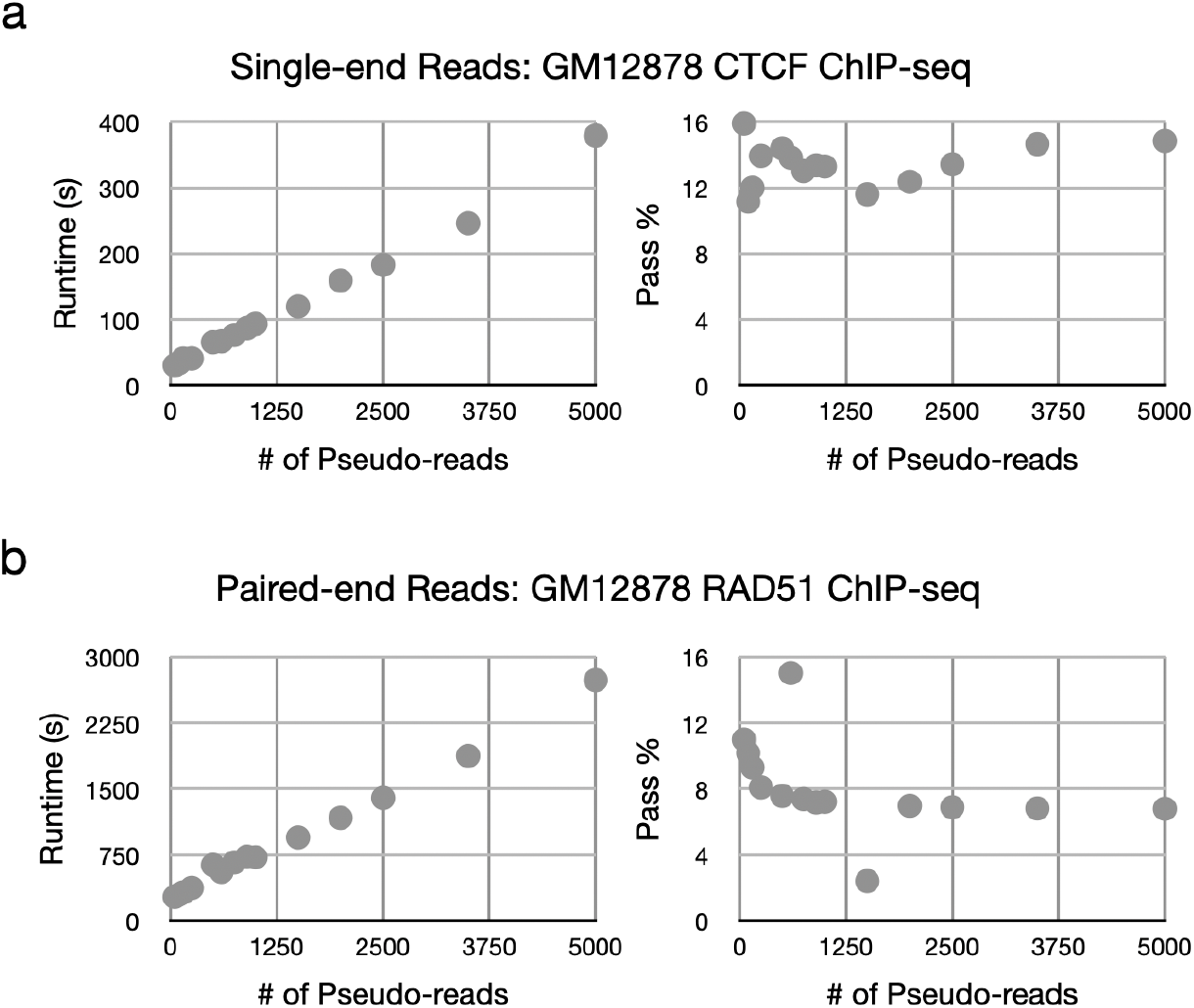
Runtime and Pass % vs. Number of Pseudo-Reads For one single-end (panel a) and one paired-end (panel b) dataset, given pseudo-reads from 50 to 5000, the runtime and pass percentage for each run. Given pseudo-reads were 50, 100, 150, 250, 500, 600, 750, 900, 1000, 1500, 2000, 2500, 2500, and 5000. All runs were done with FDR 0.1.

**Supplementary Figure S5.**
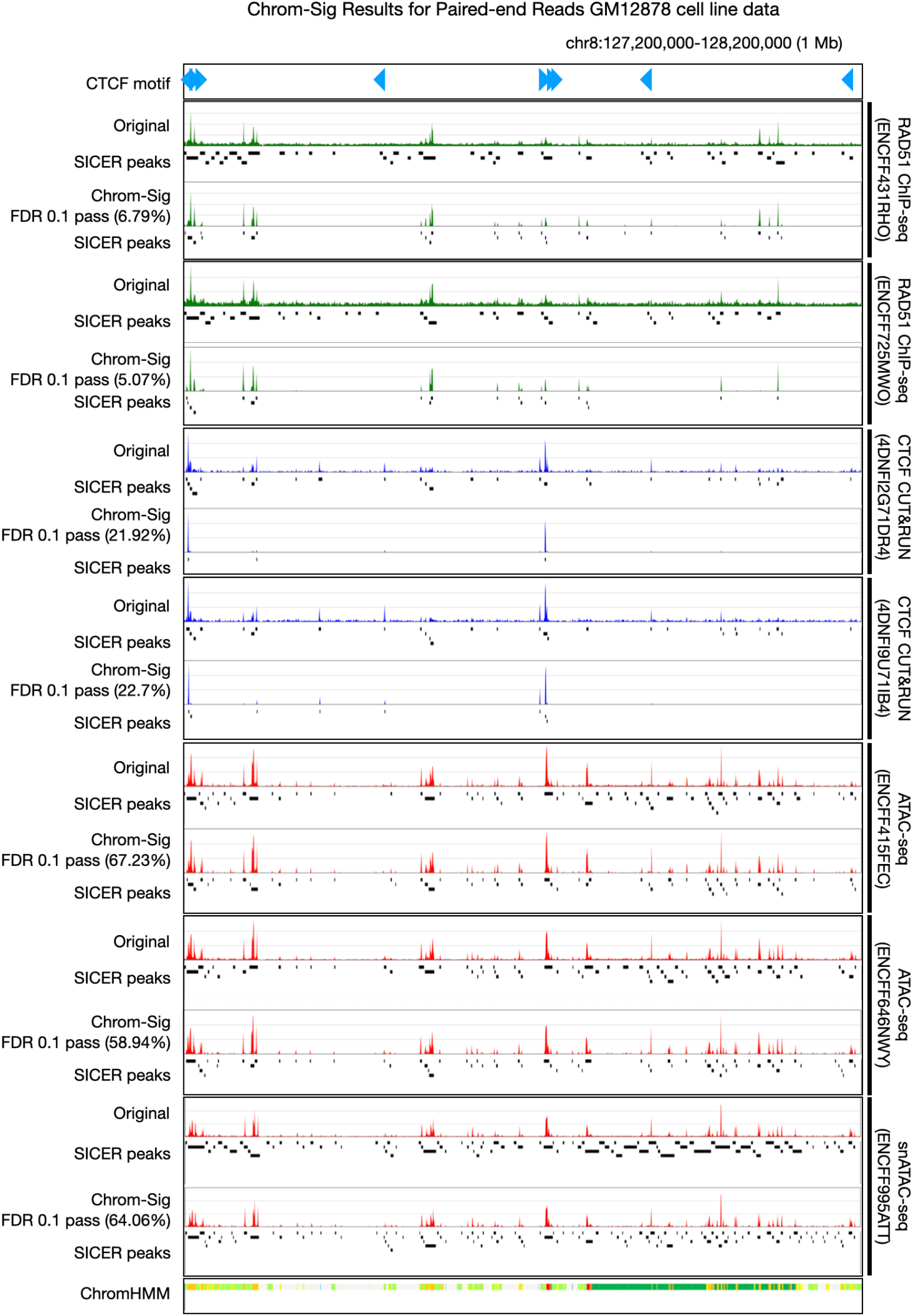
Paired End Example for GM12878 Chrom-Sig results for all paired-end datasets from GM12878 cell-line, visualized in the genome browser. CTCF Motif: CTCF binding sites with orientation. Original: Bedgraph file generated directly from input BAM/bed file. SICER peaks: Bed file result of running SICER algorithm on the original bedgraph file. Chrom-Sig FDR 0.1 pass: pass bedgraph generated from original bedgraph by Chrom-Sig (percentage refers to how many reads were retained by Chrom-Sig result from original bedgraph). SICER peaks (below Chrom-Sig FDR 0.1 pass): Bed file from SICER algorithm run on pass-pileup bed generated by Chrom-Sig. ChromHMM: Chromatin states.

**Supplementary Figure S6.**
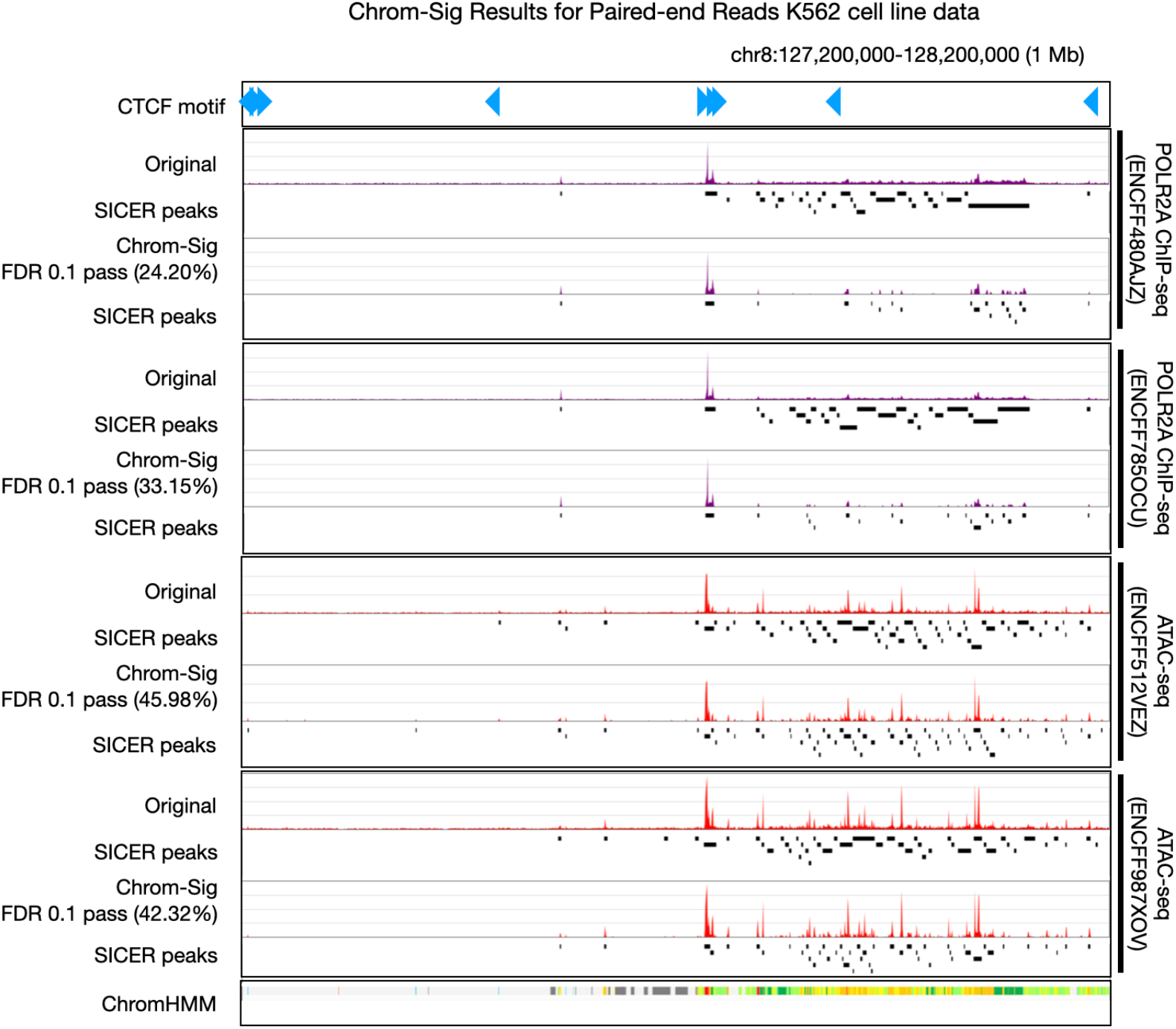
Paired End Example for K562 Chrom-Sig results for all paired-end datasets from K562 cell-line, visualized in the genome browser. CTCF Motif: CTCF binding sites with orientation. Original: Bedgraph file generated directly from input BAM/bed file. SICER peaks: Bed file result of running SICER algorithm on the original bedgraph file. Chrom-Sig FDR 0.1 pass: pass bedgraph generated from original bedgraph by Chrom-Sig (percentage refers to how many reads were retained by Chrom-Sig result from original bedgraph). SICER peaks (below Chrom-Sig FDR 0.1 pass): Bed file from SICER algorithm run on pass-pileup bed generated by Chrom-Sig. ChromHMM: Chromatin states.

**Supplementary Figure S7.**
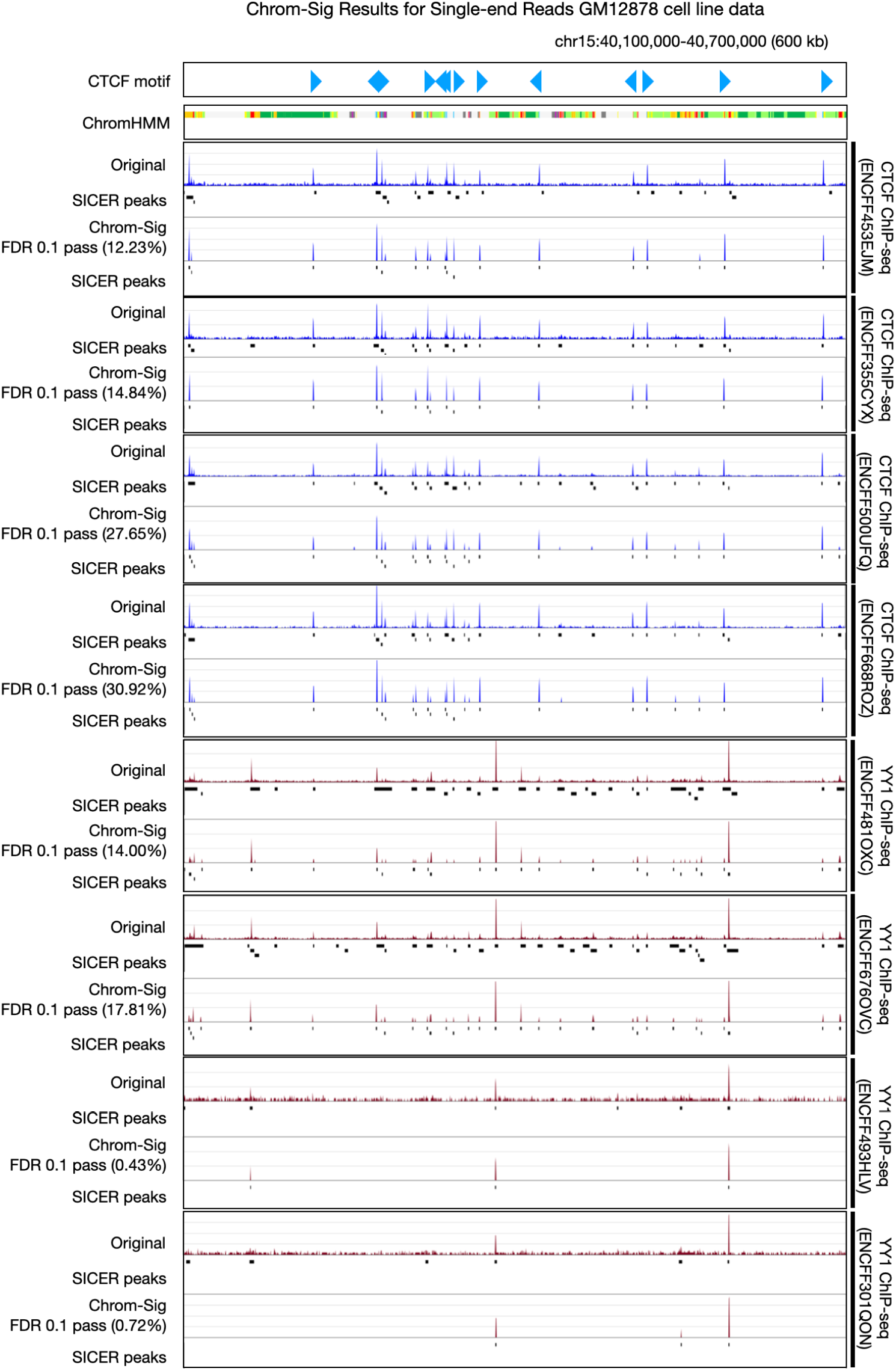
Single End Example Chrom-Sig results for all single-end datasets (all single-end data is from GM12878 cell-line), visualized in the genome browser. CTCF Motif: CTCF binding sites with orientation. Original: Bedgraph file generated directly from input BAM/bed file. SICER peaks: Bed file result of running SICER algorithm on the original bedgraph file. Chrom-Sig FDR 0.1 pass: pass bedgraph generated from original bedgraph by Chrom-Sig (percentage refers to how many reads were retained by Chrom-Sig result from original bedgraph). SICER peaks (below Chrom-Sig FDR 0.1 pass): Bed file from SICER algorithm run on pass-pileup bed generated by Chrom-Sig. ChromHMM: Chromatin states.

**Supplementary Figure S8.**
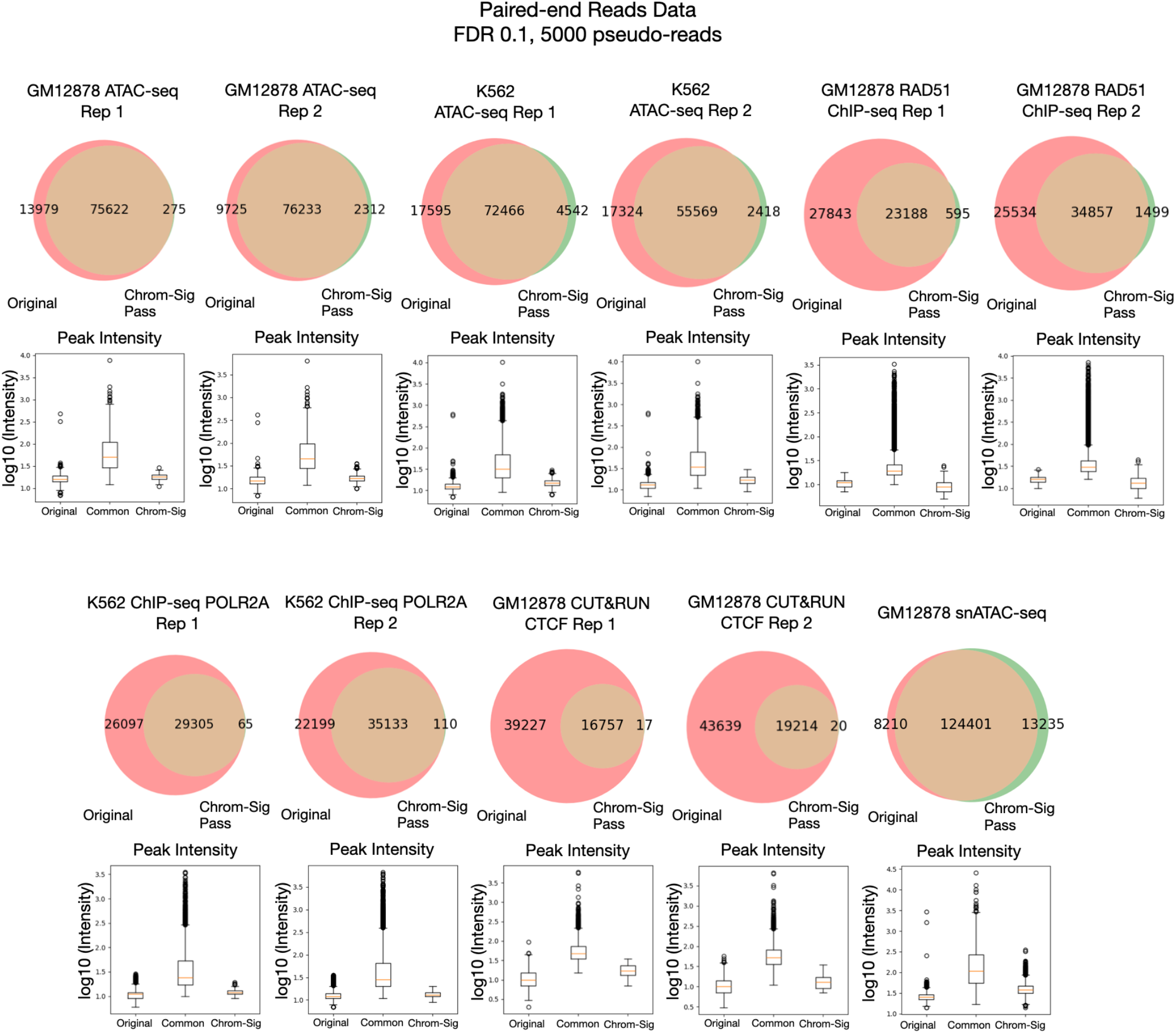
SICER Peaks in Paired End Data Before and After `Chrom-Sig` Peaks called by SICER before and after running Chrom-Sig on all paired-end datasets. Venn Diagram: Original: number of peaks called by SICER when run on bed file input (or directly generated from input). Chrom-Sig pass: number of peaks called by SICER on total (whole genome) pass-pileup bed file generated by Chrom-Sig. Box Plot: Maximum intensity of peak locations found by SICER. Intensity is determined by locating the peaks in the corresponding bedgraph file and determining the maximum using pyBedGraph. Original: Intensities of only Original SICER peaks from original bedgraph file. Chrom-Sig: Intensities of only pass-pileup SICER peaks from pass-pileup bedgraph file. Common: Intensities of SICER peaks located in both original and pass-pileup bed files. Intensity is plotted on a log10 scale. Peaks called by SICER before and after running Chrom-Sig on all single-end datasets.

**Supplementary Figure S9.**
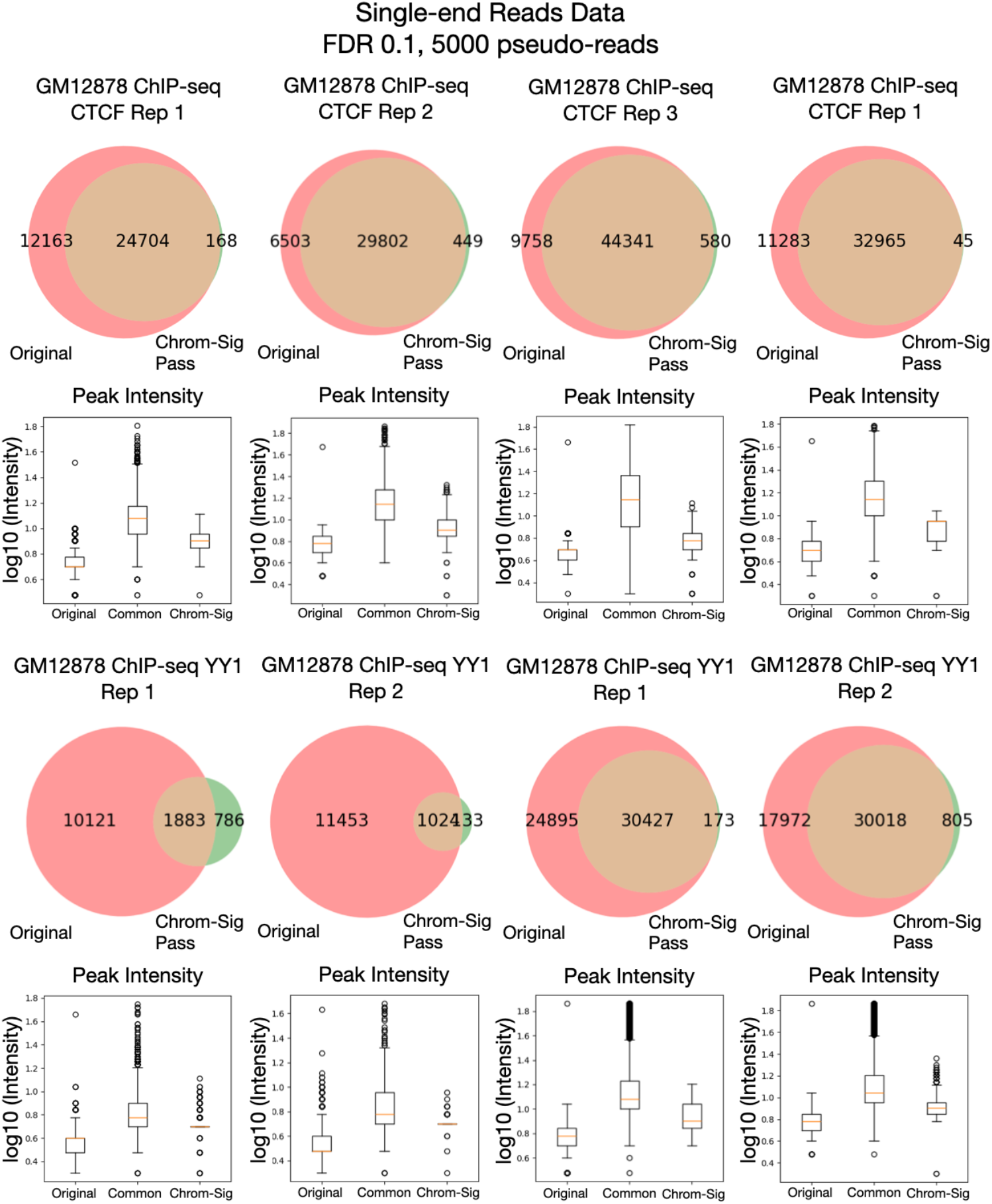
SICER Peaks in Single End Data Before and After `Chrom-Sig` Peaks called by SICER before and after running Chrom-Sig on all single-end datasets. Venn Diagram: Original: number of peaks called by SICER when run on bed file input (or directly generated from input). Chrom-Sig pass: number of peaks called by SICER on total (whole genome) pass-pileup bed file generated by Chrom-Sig. Box Plot: Maximum intensity of peak locations found by SICER. Intensity is determined by locating the peaks in the corresponding bedgraph file and determining the maximum using pyBedGraph. Original: Intensities of only Original SICER peaks from original bedgraph file. Chrom-Sig: Intensities of only pass-pileup SICER peaks from pass-pileup bedgraph file. Common: Intensities of SICER peaks located in both original and pass-pileup bed files. Intensity is plotted on a log10 scale.

**Supplementary Figure S10.**
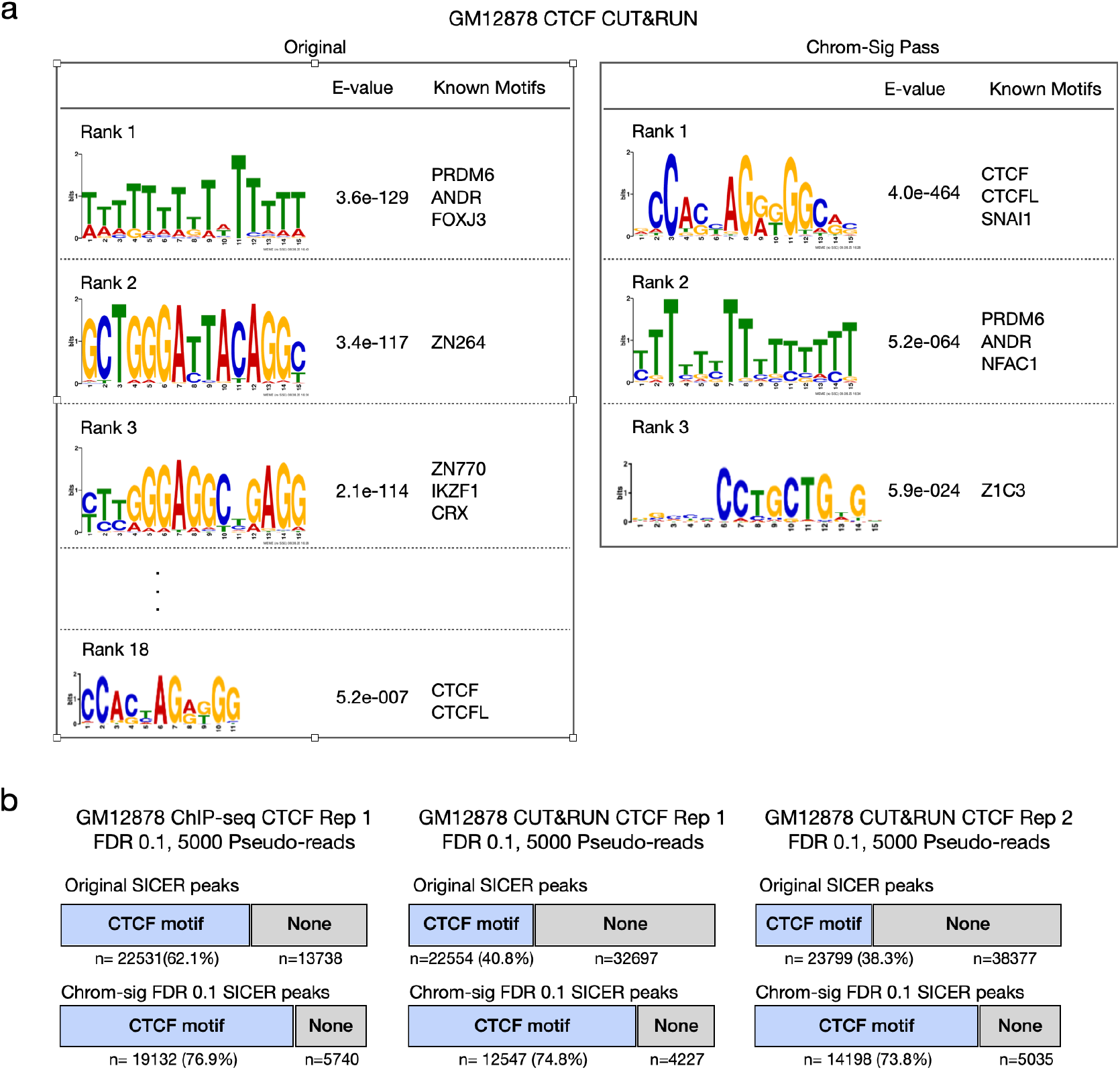
CTCF Motif Analyses a) Top enriched motifs, E-value, and matching motifs from MEME-Chip for GM12878 CUT&RUN CTCF 4DNFI2G71DR4 before and after Chrom-Sig. b) Comparison of CTCF motif precision between original data and Chrom-Sig with FDR 0.1 and 5000 pseudo-reads for GM12878 ChIP-seq CTCF ENCFF355CYX, GM12878 CUT&RUN CTCF 4DNFI2G71DR4 and 4DNFI9U71IB4.

**Supplementary Figure S11.**
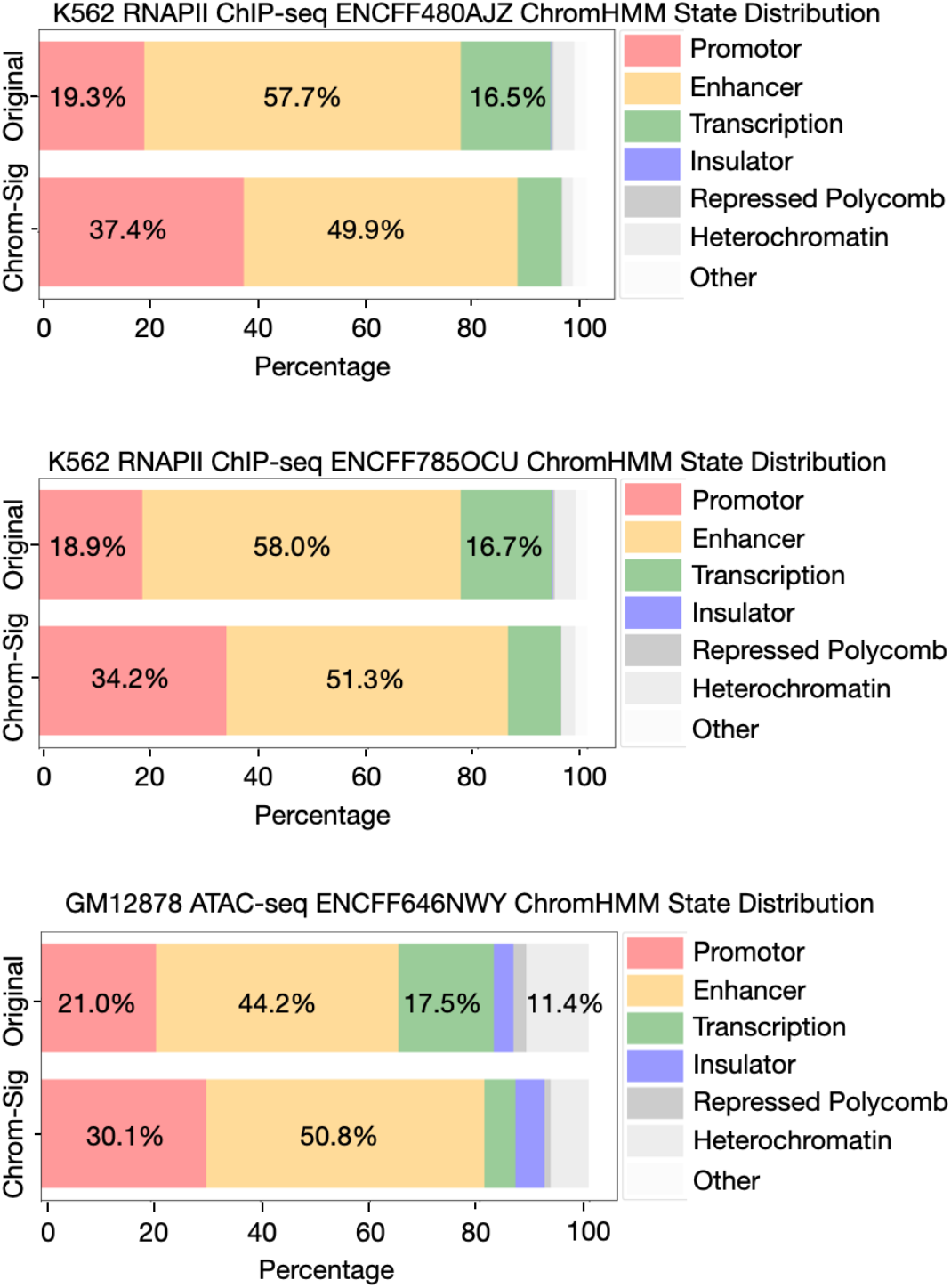
ChromHMM State Annotation Distribution Comparison of the distribution of chromHMM states between original data and Chrom-Sig with FDR 0.1 and 5000 pseudo-reads for K562 RNAPII ChIP-seq ENCFF480AJZ and ENCFF785OCU and GM12878 ATAC-seq ENCFF646NWY. The proportion of enhancer and promotor states increases when Chrom-Sig is applied to the data. Between K562 RNAPII ChIP-seq replicates there is an average of 12.3% higher distribution of enhancers and promotors (ENCFF480AJZ: 77% original vs 87.3% Chrom-Sig and ENCFF785OCU: 76.9% original vs 85.5% Chrom-Sig). In ATAC-seq data, the percentage of transcription and heterochromatin states drops from 28.9% to 12.6% after Chrom-Sig.

**Supplementary Table S1.**
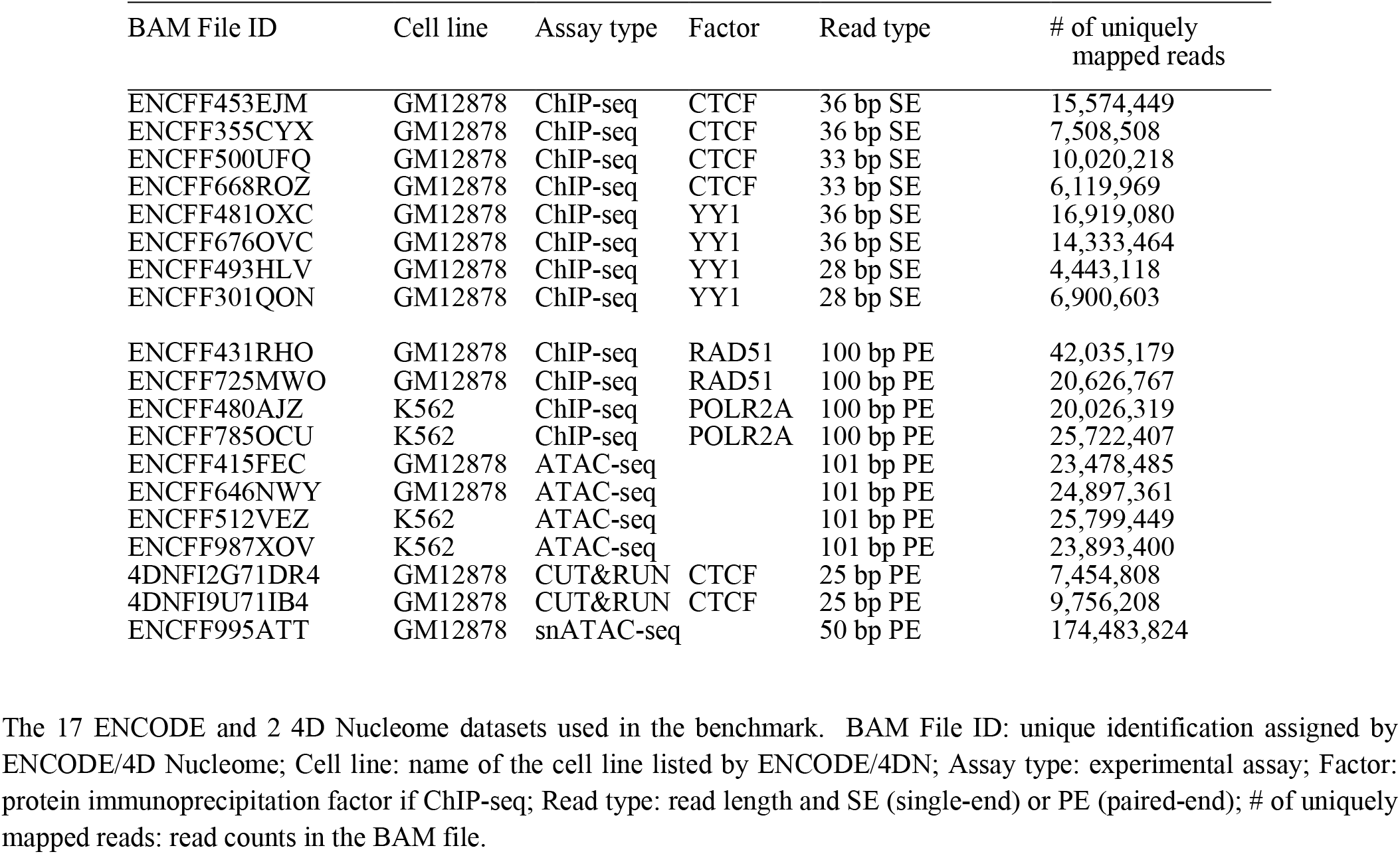
Details of 19 datasets used in the benchmark

| BAM File ID | Cell line | Assay type | Factor | Read type | # of uniquely mapped reads |
| --- | --- | --- | --- | --- | --- |
| ENCFF453EJM | GM12878 | ChIP-seq | CTCF | 36 bp SE | 15,574,449 |
| ENCFF355CYX | GM12878 | ChIP-seq | CTCF | 36 bp SE | 7,508,508 |
| ENCFF500UFQ | GM12878 | ChIP-seq | CTCF | 33 bp SE | 10,020,218 |
| ENCFF668ROZ | GM12878 | ChIP-seq | CTCF | 33 bp SE | 6,119,969 |
| ENCFF481OXC | GM12878 | ChIP-seq | YY1 | 36 bp SE | 16,919,080 |
| ENCFF676OVC | GM12878 | ChIP-seq | YY1 | 36 bp SE | 14,333,464 |
| ENCFF493HLV | GM12878 | ChIP-seq | YY1 | 28 bp SE | 4,443,118 |
| ENCFF301QON | GM12878 | ChIP-seq | YY1 | 28 bp SE | 6,900,603 |
| ENCFF431RHO | GM12878 | ChIP-seq | RAD51 | 100 bp PE | 42,035,179 |
| ENCFF725MWO | GM12878 | ChIP-seq | RAD51 | 100 bp PE | 20,626,767 |
| ENCFF480AJZ | K562 | ChIP-seq | POLR2A | 100 bp PE | 20,026,319 |
| ENCFF785OCU | K562 | ChIP-seq | POLR2A | 100 bp PE | 25,722,407 |
| ENCFF415FEC | GM12878 | ATAC-seq |  | 101 bp PE | 23,478,485 |
| ENCFF646NWY | GM12878 | ATAC-seq |  | 101 bp PE | 24,897,361 |
| ENCFF512VEZ | K562 | ATAC-seq |  | 101 bp PE | 25,799,449 |
| ENCFF987XOV | K562 | ATAC-seq |  | 101 bp PE | 23,893,400 |
| 4DNFI2G71DR4 | GM12878 | CUT&RUN | CTCF | 25 bp PE | 7,454,808 |
| 4DNFI9U71IB4 | GM12878 | CUT&RUN | CTCF | 25 bp PE | 9,756,208 |
| ENCFF995ATT | GM12878 | snATAC-seq |  | 50 bp PE | 174,483,824 |

The 17 ENCODE and 2 4D Nucleome datasets used in the benchmark. BAM File ID: unique identification assigned by ENCODE/4D Nucleome; Cell line: name of the cell line listed by ENCODE/4DN; Assay type: experimental assay; Factor: protein immunoprecipitation factor if ChIP-seq; Read type: read length and SE (single-end) or PE (paired-end); # of uniquely mapped reads: read counts in the BAM file.

**Supplementary Table S2.**
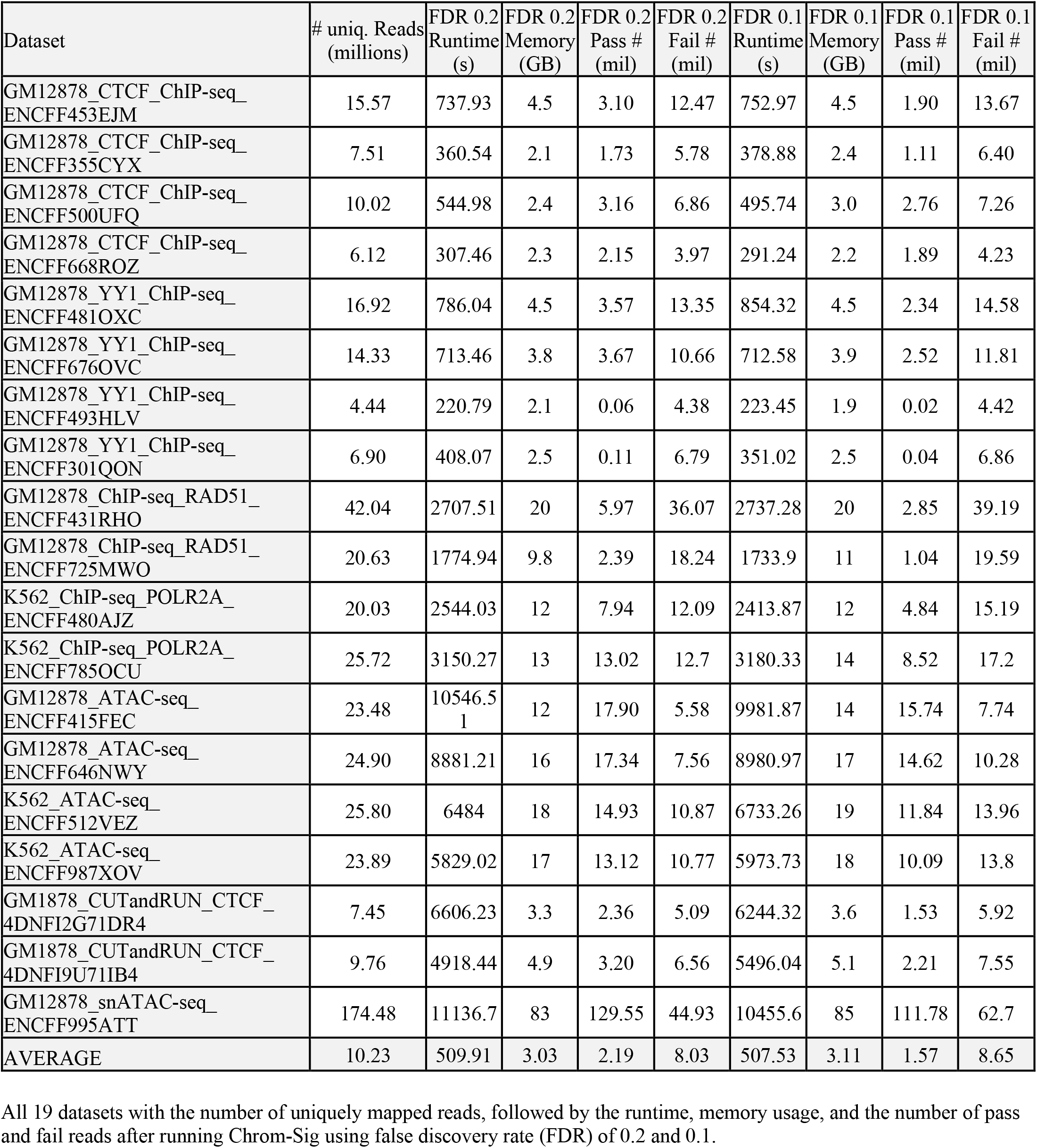
Runtime, memory, statistics for 5000 pseudo-reads

